# Novel Approach for Microbiome Analysis Using Bacterial Replication Rates and Causal Inference with Applications

**DOI:** 10.1101/2020.05.21.108514

**Authors:** Vitalii Stebliankin, Musfiqur Sazal, Camilo Valdes, Kalai Mathee, Giri Narasimhan

## Abstract

**Motivation:** Metagenomics sequencing data can be used to compute not just the relative abundance profile, but also the replication rates of every taxon in the microbiome sample. We investigate how the dynamics implied by the replication rates can be used to understand the antibiotic response in microbiomes, given the significant variation in the types of antibiotics and the types of response by different taxa. The analysis is further expanded by factoring in the resistome of the microbiomes, which can be readily profiled from the metagenomic sequence data. The fact that some antibiotics such as *β*-lactams target replicating cells makes it even more critical to use replication rates to analyze the antibiotic response.

**Results:** We introduce a novel approach for metagenomic analysis that integrates microbial community profiling, replication rate calculation, and causal structural learning to analyze the antibiotic response. First, we developed PeTRi, which involves efficient cluster computation of bacterial replication rates from metagenomic sequence data. Second, we integrate the abundance profile, replication profile, resistome profile, and environmental variables to perform causality analysis. Finally, we applied the integrated analysis to the data from an infant gut microbiome study. Conclusions from our analysis are as follows: (i) Microbes tend to lower their replication rates in response to *β*-lactams; (ii) The presence of antibiotic resistance genes combined with the causality analysis strongly suggest that genes *fosA5, oqxA, kpnF, arnA*, and *acrA* provides resistance for the taxon *K. pneumoniae*, allowing it to replicate and dominate the microbiome after the drug ticarcillin-clavulanate was administered; and (iii) Human and donor milk strongly influence the resistome of the infant gut microbiome.

**Availability:** The pipeline for calculation of bacterial replication rates using the high performance computing framework `Slurm` is available from https://biorg.cs.fiu.edu/petri/.

## 1. Introduction

Analysis of metagenomic sequencing data allows us to compute the abundance of each microbe in a microbiome [1, 2]. Recent work has made it possible to use the same data to calculate replication rates of each microbial taxon in a sample [3–6]. Replication rates provide information on how the microbial composition may be changing, thus shedding light on microbiome dynamics. This work is motivated by a desire to understand what new insight may be deduced by this new capability. As an application, we show how replication rates can provide a nuanced understanding of the characteristics of the microbiomes and their responses to antibiotic stress.

Globally, antibiotic resistance (ABR) has emerged as one of the biggest threats to public health [7]. In the US alone, annually, over 2.8 million people get severe infections from antibiotic-resistant pathogens, resulting in about 35,000 deaths [8]. Many new antibiotics are designed to target different bacterial pathways. Yet, their efficacy wanes over time as microbes develop resistance to counter the drugs. Thus, understanding the antibiotic response of a microbiome is of utmost importance.

Antibiotics such as *β*-lactams target dividing cells. In some cases, bacterial taxa are known to evade the effect of *β*-lactam exposure by halting their division temporarily [9] and then resuming their growth at the end of the treatment [10]. Some slowly growing bacteria are known to have better survival rates under antibiotic exposure [11]. Alternatively, bacterial taxa may respond to *β*-lactam exposure by producing *β*-lactamases that hydrolyze the antibiotics providing resistance. The expression of *β*-lactamases is known to be proportional to bacterial growth rates [12]. Response of division rates of bacteria to antimicrobial defines its survival model [13]. Thus, the rate of replication could serve as an indicator to track ABR and to better understand bacterial defense mechanisms against antibiotics. Therefore, the growth behavior under antibiotic exposure must be investigated in the context of a microbial community, not just with bacterial isolates [11, 14].

Surprisingly, higher replication rates do not necessarily lead to a greater abundance of the bacterial taxon [3, 4, 15]. The weak correlation may be because bacterial abundance measured is a net result of replication and death. Bacterial death is not accounted for in the replication rate computation because it represents the proportion of bacteria participating in cell division [3, 16]. By the same token, bacterial abundance, which is computed as part of standard metagenomic profiling, cannot be used as a *proxy* for replication rates.

Replication rates may be regulated by varying nutrient availability. In contrast, replication may be slowed down or halted in the presence of external stressors such as antibiotics. Since nutrients requirements and response to antibiotics vary with taxa, bacterial communities display a wide range of replication doubling times [17] and death rates [18] at any given time.

In this study, we include replication rate information to shed light on the dynamics within a microbiome. We use whole metagenome sequencing data to investigate the replication rates of each taxon in the microbiome to discern the impact of antibiotics and the development of resistance. We also use the theory of *causal inference* to study the causal relationships between the abundance and replication rates of microbial taxa, antibiotics dosages, and other clinical and environmental factors.

## 2. Approach

### 2.1. Peak-To-Trough Ratio (PTR)

Replication rates can be inferred from whole-genome DNA sequencing (mWGS) by exploiting the mechanism of bacterial cell replication [3]; circular bacterial chromosome initiate replication at the origin in both directions of the DNA strand. Thus, if bacteria are actively growing, more reads will be observed near the origin than the termination of replication. The measurement is called Peak To Trough ratio (PTR) [3] and represents the average number of replication forks per cell [16] for a given bacterial genome. Later, Brown *et al.* [4] implemented this method as an open-sourced software `bPTR` with a feature of automatic origin of replication detection.

Analysis of replication rates have been applied to several microbiome studies. For example, bacterial replication rates of some species were found to be correlated with ulcerative colitis [3], necrotizing enterocolitis (NEC) [4], and Crohn’s disease [5]. Bacterial replication rates depend on the sampling site [5, 15] and bacterial taxonomy [4], but independent of proteome composition [19]. In addition, replication rates can provide insights into antibiotic response. Bacterial taxa can lower their replication rates in response to antibiotics [19, 20] and resume their growth after cessation of the exposure [4, 20, 21]. Also, average bacterial replication were shown to be correlated with quantity of ABR genes found in a sample [21].

However, the inferences drawn about microbiomes using replication rates remain fairly simplistic. One possible reason is the complexity of computing PTR. It requires mapping the sequenced reads against a reference collection of genomes, which is hardly scalable for a large reference database, if the sequenced collection of reads has high sequence depth, or if we only have limited computing resources. A recent tool called Flint demonstrates the design of scalable algorithms for metagenomic profiling by using MapReduce, streaming, and commodity resources on the cloud [22].

In this study, we developed a scalable pipeline for calculating bacterial replication rates from metagenomic samples on a cluster environment by combining `bPTR` tool [4] with the high-performance computing approach of Flint [22]. Metagenome sequence data from a longitudinal study of sampled preterm infants were reanalyzed with bacterial replication rates information.

### 2.2. Causal Graph

Even though some studies were able to simultaneously measure the replication rates of several microorganisms [6, 23] from metagenomic samples, interactions within a microbial community were not considered in the analysis. Yet, inter-species synergies play a vital role in microbiome functioning and can create new properties not observed when studied in isolation [24]. For example, it has been shown that resident microbial communities can suppress the growth and evolution of *Escherichia coli* [25]. In this study, we aim to model complex causal relationships within microbial communities based on bacterial replication rates, microbial abundance, antibiotic-resistance genes, and miscellaneous clinical variables.

Recently, causality showed promising results to infer complex relationships (for example, colonization pattern) among different entities of a microbiome [26, 27]. Causal inference technique is useful in the classification of pathogenic and beneficial bacteria just using microbial taxa abundance information [28]. The first step of causal inference is to establish relationships among the entities, and the idea of causal structure is as follows.

Formally, we define *causal structures* (CS) (or *causal Bayesian networks*) as a class of Probabilistic Graphical Models (*PGMs*) [29, 30] where each node represents one of *n* random variable from a set, **X** = {*X_i_*, *i* = 1,…,*n*}, and each edge represents a direct causal relationship. These structures are represented as a graph *G* = (*V, E*), where each vertex in *V* represents a random variable from **X**, and *E* is the set of edges. Although undirected edges are used in cases where the direction cannot be reliably determined or when both directions appear to be valid, the graph *G* is often “manipulated” to be a Directed Acyclic Graph (DAG). Each random variable *X_i_* has an associated probability distribution. A directed edge in *E* between two vertices represents direct stochastic dependencies. Therefore, if there is no edge connecting two vertices, the corresponding variables are either marginally independent or conditionally independent (conditional on the rest of the variables, or some subset thereof). The “local” probability distribution of a variable *X_i_* depends only on itself and its parents (i.e., the vertices with directed edges into the node *X_i_*); the “global” probability distribution, *P* (**X**) is the product of all local probabilities, i.e., a joint distribution [31], given by

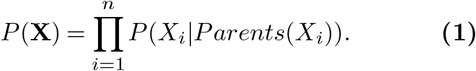

Note that the equation is simpler when the causal structure is sparser. Thus, an important step in our pipeline is to identify all independent pairs of random variables. More importantly, we also identify as many conditionally independent pairs as possible since these represent indirect or non-causal relationships.

All local structures in a causal structure can be classified into three sub-categories: *chains*, *forks*, and *colliders*. In a chain, two variables *X* and *Y* are conditionally independent given *Z*, if there is only one unidirectional path between *X* and *Y*, and *Z* is the set of variables that intercepts that path. In a fork, variable *Z* is a “common cause” for variables *X* and *Y*; this happens when there is no directed path between *X* and *Y*, and they are independent conditional on *Z*. Finally, variable *Z* is a “collider” node between *X* and *Y*, if it is the “common-effect”. In a collider, as in the fork, there is no directed path between *X* and *Y*. However, the difference is that *X* and *Y* are unconditionally independent, but become dependent when conditioned on *Z* and any descendants of *Z*.

In general, causal models can be very complex. A pair of variables can be connected through multiple chains, forks, and colliders, making it non-trivial to determine the dependency between two arbitrary variables. *Directional separation* (or, just *d-separation*) is a useful concept in this context [32] because covariance terms corresponding to *d*-separated variables are equal to 0. In a directed graph, *G*, two vertices *x* and *y* are *d*-connected if and only if *G* has a collider-free path connecting *x* and *y*. More generally, if *X, Y* and *Z* are disjoint sets of vertices, then *X* and *Y* are *d*-connected by *Z* if and only if *G* has a path *P* between some vertex in *X* and some vertex in *Y* such that for every collider *C* on *P*, either *C* or a descendant *C* is in *Z*, and no non-collider on *P* is in *Z*. *X* and *Y* are *d*-separated by *Z* in *G* if and only if they are not *d*-connected by *Z* in *G*. The concept of *d*-separation allows for more edges to be eliminated in a causal structure.

Meaningful causal relationships inferred from causal Bayesian networks constructed using principles of *d*-separation are more informative than association analysis. A directed edge from *X* to *Y* in such networks suggests that changing X will imply a change in the value of Y, if all other variables are held unchanged [33]. Therefore, with PGMs, it is possible to infer the causal impact of one variable on another, assuming that there are no hidden confounders. Even though the assumption does not hold true in reality, approximations provided by the model give valuable insights into causal relationships in a multivariate data set.

## 3. Methods

In this paper, we propose a new approach to study antibiotic resistance from microbiome sequence data. Figure 1 shows an overview of the proposed three-pronged approach. First, Figure 1 (A) presents a pipeline to efficiently calculate bacterial replication rates and relative abundance values using a high-performance computing environment. Second, (B) shows the framework to quantify ABR genes from the sequencing data. Third, we compute a causal network, where the allowed edges are as shown in the skeleton network in (C).

**Fig. 1.**
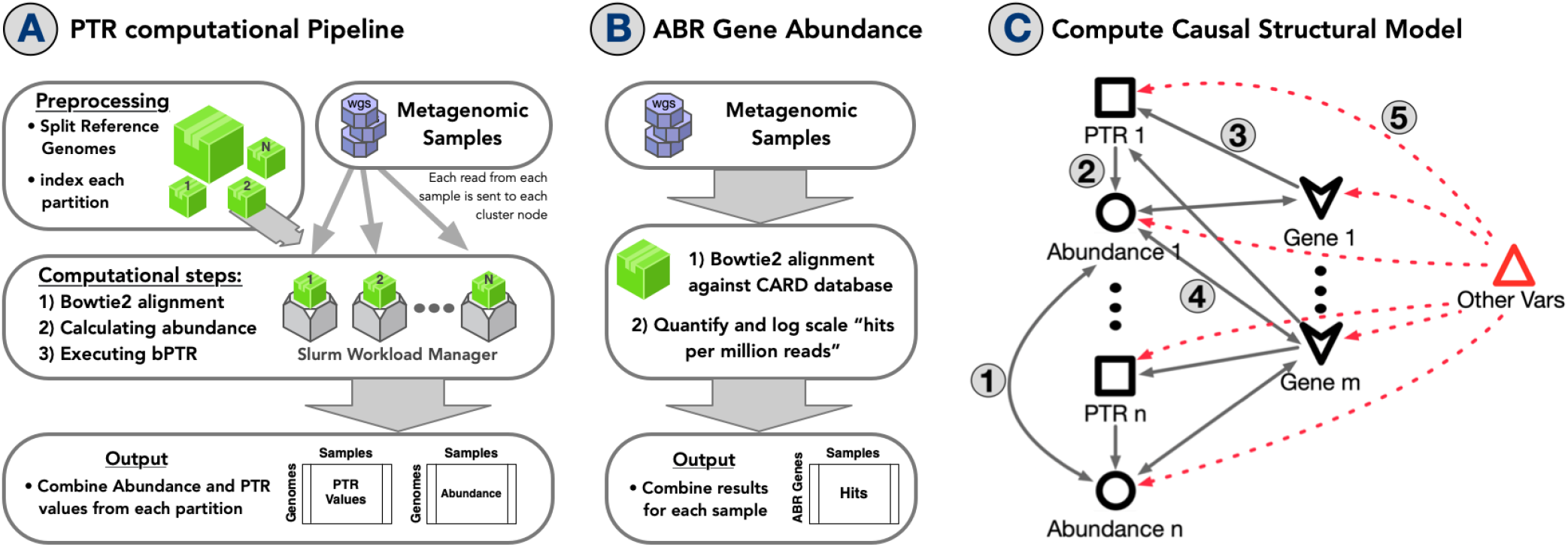
Proposed approach to study relationships between replication and antimicrobial resistance **A.** PeTRi pipeline for replication rates computation; **B** Pipeline for quantifying ABR genes; **C**. Model for causal structure where allowed edges are shown numbered 1-5. Different shapes are used for different types of nodes.

### 3.1. PeTRi: A PTR Calculation Pipeline

To calculate bacterial replication rates for a large number of samples and reference genomes, we created a pipeline called PeTRi that runs on a high-performance multiprocessor computing environment with the Slurm resource management system [34]. The pipeline consists of three steps — invoking the `bowtie2` alignment software to align the sequenced reads [35], calculating the abundance of each taxon in the sample using Flint [22], and calculating growth rates of each taxon using the `bPTR` [4] software.

All reads were mapped by `bowtie2` to a reference genome collection containing 30,382 complete bacterial, archaeal, and viral genome sequences from the RefSeq repository [36]. Genomes were downloaded using the API offered by `Kraken 2` [37].

To utilize Flint in an optimal manner, we split the database into 64 roughly equal-sized pieces and created `bowtie2` indexes as described previously [22]. As part of the preprocessing, the indexes are loaded onto the 64 different computational nodes in the cluster. Thus, each read is sent to all the 64 units, which run with their own local index of a subset of genomes from the repository.

As part of the first step in the processing pipeline, `bowtie2` is used in each processor to align every read to its local index partition. Next, the alignment file is analyzed to calculate the average coverage of each genome using the following formula:

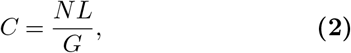

where *C* is the average coverage, *N* is the total number of reads that align to the given genome sequence, *L* is the average length of a metagenomic read, and *G* is the genome sequence length. Next, we compute the relative abundance of each genomic taxon by normalizing average coverage values to add up to 1. We filter out the list of genomes with average coverage less than 5, which is the minimum coverage requirement for `bPTR`. Finally, we run the `bPTR` software on the set of genomes that survived the aforementioned filtering step. At the end of the processing step, for each sample, we obtain the PTR values for each genome along with the corresponding relative abundance values. The stringent filtering step is required to remove false positives from the mapping step where reads may get mapped to multiple closely related microbial strains.

Finally, we compute species-level PTR values using the following formula:

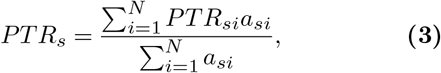

where *PTR_s_* is the PTR value for species *s* with *N* strains; *PTR_si_* is the PTR value of strain *i* of species *s*; and *a_si_* is the relative abundance of strain *i* of species *s*. Such an aggregation gives us the average PTR value for a given species *s*, and reduces the variance between values, providing more robust statistical results. Note that weighting the PTR with abundance eliminates anomalies that arise from extremely high PTR values of strains with low abundance. Additionally, it remains stable even if strains evolve over time.

The PeTRi pipeline described above has many advantages over existing approaches. Our method is readily scalable. It can deal with a large number of samples, and associated sequencing reads because of the use of a cluster. In addition, our method can be scaled to deal with a huge reference genome collections. Splitting the indexes for the reference genomes database into smaller manageable pieces provides an efficient framework for the scalable computation of PTR values. The associated computation times are shorter, and the costs are meager if they are performed on a cloud service such as AWS [38]. Using more extensive genome collections allows us to measure PTR values for many more strains that may be present in the samples than currently possible with other tools, which use much smaller reference genome collections, as discussed below. Existing tools for computing PTR values such as `GRiD` [6] use curated reference genome collections with at most one genome per species. This is likely to limit the sensitivity of the PTR profile that is computed since strains not represented in the reference collection may have very different replication rates than the representatives. A recently developed tool called `SMEG` [23] deals with strain variations by finding representative reference sequence. However, we were unable to parallelize `SMEG` in a natural way, making it computationally slower than the tools we have used. Other tools for computing PTR values such as `iRep` [4] and `Demic` [5] use draft-quality metagenome-assembled genomes. Consequently, the computed PTR values are really coarse averages across multiple strains since reconstructed genomes often have to be dropped because of insufficient coverage or contain mosaics of multiple strains. Furthermore, metagenomic assembly and binning tasks are computationally intensive.

### 3.2. Analyses of ABR Genes

DNA sequences for a list of 2624 antibiotic-resistance genes were downloaded from the Comprehensive Antibiotic Resistance Database (CARD) version 3.0.7 [39]. Metagenomic reads were then aligned against these sequences using `bowtie2`, allowing us to calculate “counts per million reads” for each ABR gene. Once normalized, these measures are used as a proxy for the antibiotic resistance “potential” of the metagenomic sample. Note that this potential is a substitute for the expression of the ABR genes in the sample. The CARD database also has information on the bacterial taxon from which the ABR gene sequence was isolated. It is important to note that it is not possible to determine (either from the metagenomic or metatranscriptomics sequence data) the specific bacterial taxon that carries the ABR gene. However, the annotation from the CARD database provides a useful prediction.

### 3.3. Causal Inference

To infer causal relationships between the variables of interest, we used the PC-stable algorithm [40] using bnlearn [41] R package. The multi-variate random variables used for our causal analysis include: (a) relative abundance of a taxon, (b) PTR value of a taxon, (c) dosage of maternal antibiotics administered, (d) dosage of infant antibiotics administered, (e) log-scaled abundance of a antibiotic resistance gene, and (f) miscellaneous environmental factors (e.g., caffeine, mother’s milk, donor milk, infant birth weight, iron supplement, etc.). Note that each node of the causal network represents a multi-dimensional variable, one dimension for each sample.

To ensure biologically relevant causal structures, we restricted the set of allowed edges. Similar restrictions were introduced in Lugo-Martinez *et al.* [42]. As shown in Figure 1-C, the following types of edges are allowed:

**Type 1:** Taxon abundance ↔ Taxon abundance
**Type 2:** PTR → abundance (of the same taxon)
**Type 3:** ABR gene → PTR
**Type 4:** Taxon abundance ↔ ABR gene
**Type 5:** Other vars → {Abundance, PTR, ABR gene}

Edges of Type 1 between abundance values of different taxa represent microbial interactions such as co-operation or competition [43]. Edges of Type 2 suggest that a change in replication rate may change the abundance of that taxon. This paper has evidence suggesting that the presence of an ABR gene may promote the replication of the taxon that carries it (Type 3 edge). Type 4 edges suggest that higher abundance implies increased copies of the ABR gene in that sample. This may mean that increased expression may confer resistance. Finally, clinical and environmental variables are independent variables and impact all aspects of the microbiome (Type 5 edges).

All edges were filtered using a bootstrap threshold of 0.5. To keep the resulting models simple and sparse, we constructed a model using only the most abundant major bacterial species and AMR genes present in at least 5 % of the samples from the study.

### 3.4. Statistical Discriminant Analysis

To determine if antibiotics significantly affected the microbial resistome, we performed Permutational Multivariate Analysis of Variance (PERMANOVA) tests [44] using the vegan R package [45] using Jaccard similarity. PERMANOVA is a non-parametric test used to compare two (or more) groups of objects and test the null hypothesis that the centroids and dispersion of the groups using the distance measure are equivalent for the groups.

To visualize the differences between the two classes, we used Principal Coordinates Analysis (PCoA). PCoA is a dimensionality-reduction technique that embeds the objects in a low dimensional space so that distances are as faithfully reflected in the new space. The PCoA models were trained using ape R package [46].

To identify the antibiotic-resistance genes that help to discriminate between the different antibiotics, we applied the Partial least squares discriminant analysis (PLS-DA) technique for feature selection using the `MixOmics` R package [47]. The PLS-DA is a supervised machine learning tool that finds the direction (i.e., principal component) that maximizes the separation among the classes. To select the set of features that contribute to the class separation the most, we used 10-fold cross-validation and 10 repeats with tune.splsda. Statistical significance was computed using the Mann-Whitney U tests.

### 3.5. Hardware

PeTRi pipeline (Section 3.1) was executed on High-Performance Computational (HPC) resources at FIU Instructional and Research Computing Center [48]. The job submission was handled by Slurm Workload Manager version 19.05.3-2. The cluster contains 1500 Intel-based cores with High Memory Nodes ranging from 32 to 384 GB per node. All jobs were submitted to computational nodes with CentOS v.7 operating system. To be able to execute several jobs within the same computational node, we used only eight threads per one process.

Other computational steps (Sections 3.2-3.4) were performed on a server machine at FIU, `castalia`, with 792GB of RAM, and 48 Intel Xeon processors.

## 4. Results and Discussion

We applied our pipeline to the premature infant gut microbiome data set (BioProject ID: PRJNA301903) [49] and calculated bacterial replication rates. The data are from 401 stool samples from 84 premature infants [49]. All but two infants received antibiotic therapy within the first 24 hours. Forty-nine of the infants received additional antibiotic therapies (“Antibiotic” cohort) between 1–10 weeks of life. Remaining 35 formed the “Control” group. Each therapy consisted of one or more antibiotics (ampicillin (AMP), meropenem (MEM), ticarcillin-clavulanate (TIM), gentamicin (GEN), vancomycin (VAN)). Some treatments were occasionally mixed with Cefazolin (CFZ), Trimethoprim-Sulfamathoxazole (SXT), Clindamycin (CLI), or Cefotaxime (CTX). Stool specimens were collected at roughly 2-6 samples per infant at multiple time points between 6-156 days of life.

### 4.1. Does PTR Correlate with Abundance?

We first argue that contrary to expectations, replication rates does not positively correlate with relative abundance. In other words, increased replication does not lead to higher relative abundance values. Fig. 2-A shows the Spearman correlation between PTR and relative abundance values of the major bacterial species present in the dataset. This is also consistent with the results of Brown et al. [4]. For eight out of 25 taxa, the correlation is negative, while three species (*Bifidobacterium longum, Enterococcus faecium* and *K. michiganensis*) have a positive correlation in both premature and healthy infants.

**Fig. 2.**
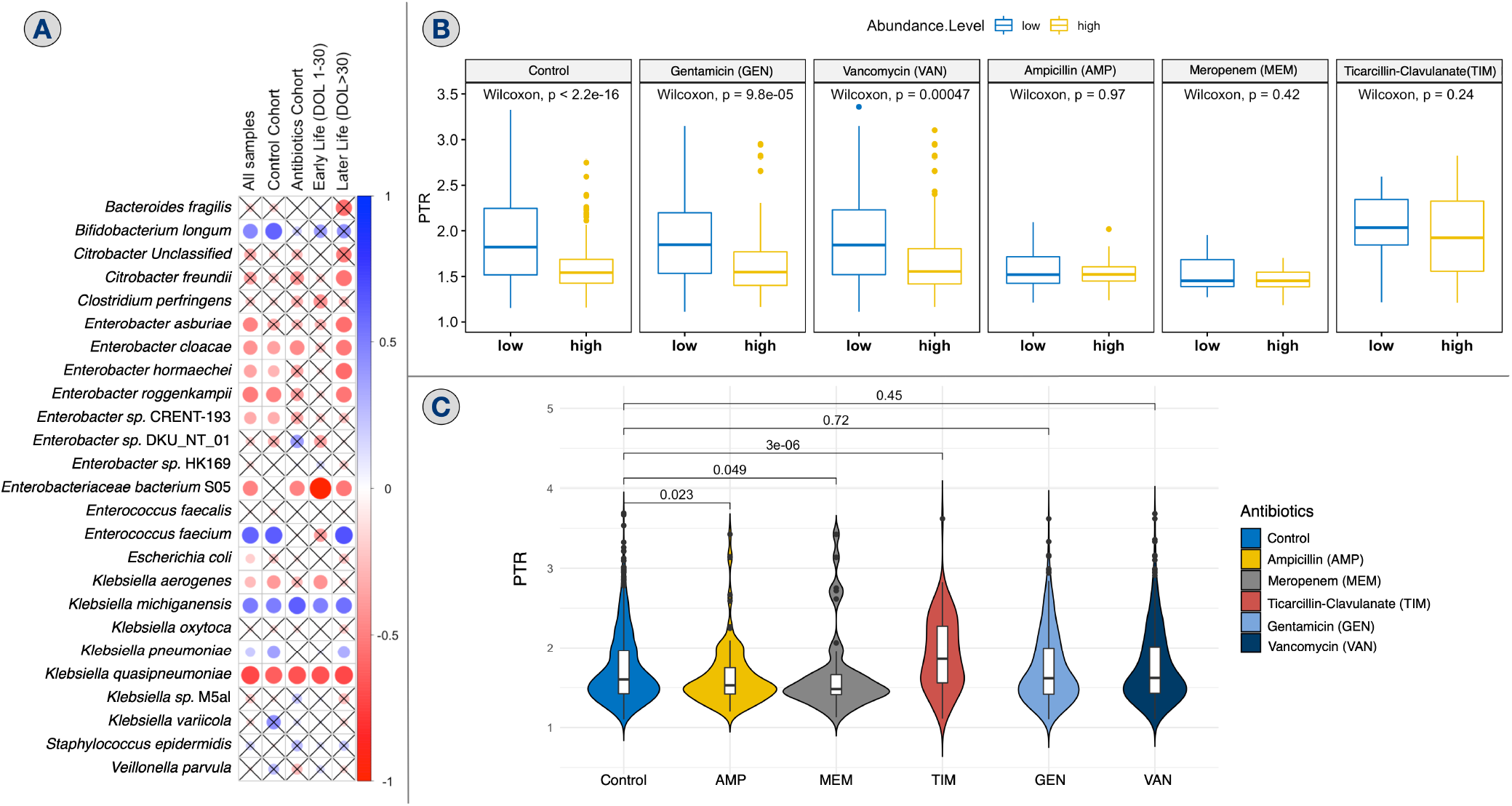
**A.** Spearman correlation coefficients between PTR and relative abundance values of the same taxon as part of different cohorts. Species that were present in less than 20 samples were not included in this analysis. Color of the filled circles signifies the sign of the correlation (red for positive and blue for negative), while the size of the filled circles represents the absolute value of the correlation coefficient. Crossed entries represent correlations that are not significant (p-value > 0.05) **B.** Box and whisker plots of PTR values of “low” and “high” abundance bacterial taxa, broken down by the type of antibiotics administered within 48 hours prior to sampling. The “low” group includes taxa with the lowest 25% relative abundance values (values < 0.1%), while “high” group consist of highest 25% abundant taxa (values >4%) **C** Violin plots showing distribution of PTR values in control infants and infants who recently received antibiotic treatment. Mean PTR in each treatment group was compared to control samples using the Mann–Whitney U test (p-values are shown in the figure).

Negative correlations between PTR and abundance values were for most part consistent in the two cohorts – the “Control” cohort (infants with no additional antibiotic treatment) and the “Antibiotics” cohort (infants with additional treatment). It was also true when we sliced the data set differently, i.e., for the “Early Life” samples (first 30 days of life) as well as for “Later Life” samples (after first 30 days of life). The only inconsistencies seen were with the taxa, *Enterococcus faecium* spp. and *Enterobacter* DKU_NT_01, and with the subset of samples immediately after the administration of the *β*-lactam antibiotics (discussed below).

There are more cases of negative correlation (than positive) between the replication rates of taxa and their abundance. This finding implies that exhaustion of essential nutrient leads to the bacterial taxa reaching a stationary phase [50]. Figure 2-B shows PTR values of the taxa that ranked among the highest 25% (high) and lowest 25% (low) relative abundance taxa for each type of recently administered antibiotics (within 48 hours prior to sampling). Replication rates of taxa with high abundance were significantly lower than that of less abundant bacteria in the “Control” group, as well as in the cohorts that were treated with gentamicin and vancomycin.

Higher abundance did not imply lower replication when *β*-lactams were administered. No significant difference existed between PTR of “high” and “low” abundant species with the administration of *β*-lactams - MEM, AMP, and TIM. Disruption of replication is explained by the fact that *β*-lactams target dividing cells [51].

### 4.2. How are the PTR values distributed?

Distribution of replication rates values for each class of recently administered antibiotics is shown in Fig. 2-C. There was no significant difference between the mean PTR for the control group compared to those that were recently administered GEN or VAN. After the administration of AMP or MEM, the average PTR is lower compared to control. As mentioned earlier, this appears to be in response to the *β*-lactam antibiotics [10]. The lowest replication values were observed when MEM was administered, perhaps because degradation by *β*-lactamases appears to be least effective against this antibiotic [52]. Surprisingly, even though TIM belongs to *β*-lactam class, the average PTR is close to 2 after it was administered indicating that the dominant taxa were not in stationary phase after the drug exposure.

### 4.3. Responses to *β*-lactams vis-à-vis the resistome

If the reason for the increased replication rates of some taxa after the administration of TIM is their resistome, then it makes sense to investigate the resistome of the microbiome after the administration of TIM and the other *β*-lactams. PERMANOVA analysis shows that the microbial composition as well as the resistome of the infant microbiomes is significantly different for the control group and the cohorts treated with TIM and AMP/MEM (p-value <0.01; Supplementary file `supplS1.xlsx` - S1-A). In Fig. 3-A the differences in microbial abundance (left plot) and resistome (right plot) are shown using PCoA. Samples in the “TIM” cohort clustered more compactly suggesting a more consistent response. We note that points corresponding to the infants in the “Control” group occur closer to “TIM” cluster than the points from the “AMP/MEM” cohort. This may be because even the infants in the “Control” cohort were exposed to antibiotics during their first few days of life, possibly shaping their resistome, which gets expressed differently based on the exposure. On the other hand, the “TIM” and “AMP/MEM” cohorts favored the survival of a microbiome with different resistomes.

**Fig. 3.**
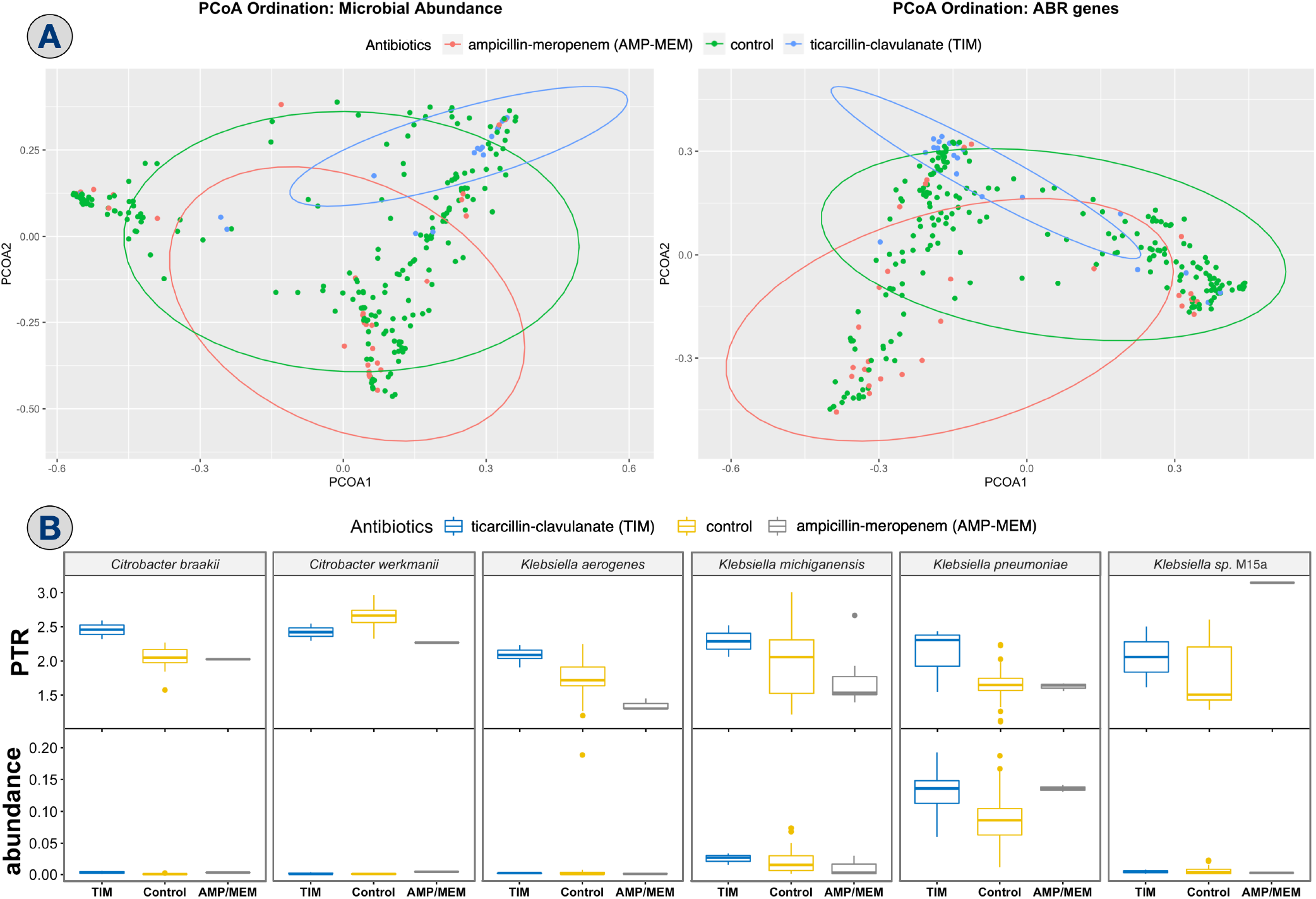
**A** Principal coordinates analysis (PCoA) ordination based on microbial relative abundance (left) and log2-transformed “hits per million reads” of ABR genes (right). Samples represent points colored by the type of antibiotics administered within the 48 h prior to sampling. **B** Box-and-Whisker Plots of PTR (top row) and abundance (bottom row) values of bacterial taxa that were in active growth after administration of ticarcillin-clavulanate

### 4.4. Replicating taxa after antibiotics

The taxa with the highest PTR values after administration of TIM were *Citrobacter braakii*, *Citrobacter werkmanii*, *K. pneumoniae*, *K. aerogenes*, *K. michiganensis*, and *Klebsiella* sp. M5al. The average PTR and abundance values of taxa in the active growth phase (PTR > 2) after TIM administration are shown in Figure 3-B. The dominant species in the TIM cohort with a mean relative abundance of 16% was *K. pneumoniae*. Despite its high abundance, this species was actively growing with high PTR values, suggesting a tendency for severe dysbiosis and pathogenic dominance by this taxon. This is further evidenced by the low abundance of other potentially pathogenic species with high PTR values such as *K. aerogenes* and *Klebsiella* sp. M5al. This finding may explain why there is a high abundance of pathobionts in premature infants, an unintended consequence of antibiotic treatment [49, 53]. Our findings suggest that the use of TIM for premature infants should be discontinued.

### 4.5. Identifying ABR genes of interest

PLS-DA and Mann–Whitney U tests were used to identify specific ABR genes that contribute the most to differentiate the resistomes of the TIM from AMP/MEM cohorts. The model was trained to differentiate two cohorts: infants recently administered AMP/MEM and the TIM cohorts. To avoid overfitting the PLS-DA model [54], a 10-fold cross-validation was employed resulting in a low error rate of 0.12. We identified 320 ABR genes that contributed the most to differentiating the cohorts. Using the Mann–Whitney U tests and Benjamini-Hochberg p-value correction, the genes of interest were reduced from 320 to 46 (see Suppl. file `supplS1.xlsx` - S1.B)

Of the 46 genes idenitfied in the analysis above, 30 were positively correlated with average PTR values of the TIM cohort (Table 1), of which 26 were associated with *K. pneumoniae* (shown in bold font), meaning that they were either originally isolated from it [62] or was present in at least 30% of all whole-genome shotgun assemblies available frmo NCBI for *K. pneumoniae*, as annotated in CARD [39].

**Table 1.** ABR genes that positively correlate with replication rates and TIM treatment.

| ABR Gene Family | Genes correlated with average PTR * |
| --- | --- |
| *K. pneumoniae* OKP chromosomal β-lactamase | ***bla*OKP.A.3, *bla*OKP.B.1, *bla*OKP.B.2, *bla*OKP.B.3, *bla*OKP.B.4, *bla*OKP.B.5, *bla*OKP.B.6, *bla*OKP.B.7, *bla*OKP.B.8, *bla*OKP.B.9, *bla*OKP.B.10, *bla*OKP.B.11, *bla*OKP.B.13, *bla*OKP.B.17, *bla*OKP.B.18, *bla*OKP.B.19, *bla*OKP.B.20** |
| Major facilitator superfamily (MFS) antibiotic efflux pump | ***kpnF*** *, *K. pneumoniae kpnG*, *K. pneumoniae kpnE* |
| Resistance-nodulation-cell division (RND) antibiotic efflux pump | ***oqxA*** *, ***oqxB***, *mexB*, *smeE*, *K. pneumoniae acrA* *- *tolC* |
| Other | *arnA* *, *fosA5* *, ***fosA6***, *K. pneumoniae ompK37* |

\*Genes marked with asterisks are predicted to be growth-enhancing based on causality analysis; In bold we show genes associated with *K. pneumoniae*.

*K. pneumoniae* was the most dominant species in the TIM cohort. The 26 ABR genes were associated with a resistance response to TIM, which correlates with PTR values. All the genes identified in this study as significant confer multidrug resistance to *K. pneumoniae*. The *K. pneumoniae* clinical isolates form three phylogenetic groups KP-I, KP-II, and KP-III [62, 63]. *K. pneumoniae* are intrinsically resistant to many *β*-lactams due to expression of a chromosomally encoded *β*-lactamase constitutively conferring resistance to ampicillin, amoxicillin, carbenicillin, and ticarcillin, but not to extended-spectrum *β*-lactams. Three groups of *bla* genes also evolved parallelly *bla*_SHV_, *bla*_OKP_, and *bla*_LEN_. The *bla*_OKP_ family was found in group KP-II [62], and *bla*_OKP-A_ and *bla*_OKP-B_ are two variants [62]. *K. pneumoniae* strain harboring *bla*_OKP_ are common in acute care settings and serves as a reservoir for this gene [64, 65]. *K. pneumoniae* harbors many genes that confers multi-drug resistance, such as those genes encoding efflux pumps (*oqxAB, kpnEF, kpnGH, acrA-tolC*), outermembrane porin protein (*ompK37*), *acrA-tolC* (confers colistin resistance), *fosA* (plasmid-encoded gene that confers fosfomycin resistance). The *K. pneumoniae kpnEF* system confers resistance to cefepime, ceftriaxone, colistin, erythromycin, rifampin, tetracycline, and streptomycin [66]. The *kpnGH* system confers resistance to azithromycin, ceftazidime, ciprofloxacin, ertapenem, erythromycin, gentamicin, imipenem, ticarcillin, norfloxacin, polymyxin-B, piperacillin, spectinomycin, tobramycin and streptomycin [67]. The *oqxAB* genes are also common in clinical isolates of *Enterobacteriaceae* and *K. pneumoniae*, conferring low to intermediated resistance to quinoxalines, quinolones tigecycline, and nitrofurantoin [68]. Loss in *ompK37* in *K. pneumoniae* confers resistance to cefoxitin and expanded-spectrum cephalosporins [69–71]. The *fosA* encoding fosfomycin modifying enzyme of plasmid origin are originally identified in *Serratia marcesens* [72, 73].

To understand which associations are causal in nature, we discuss the causality inference approach next.

### 4.6. Identifying causal relationships

The goal of this section is to better understand antibiotic resistance response exhibited in a cohort of samples by identifying which associations or correlations are causal. The causal inferencing was performed with the same data, resulting in a graph where the nodes correspond to bacterial (relative) abundance, bacterial replication rates (PTR), abundance of ABR genes, antibiotics dosage (infant and mother), and miscellaneous environmental factors (e.g., dietary information, infant age (premenstrual and chronological), and birth-weight). Figure 4 represents the resulting causal network after removing nodes without any outgoing or incoming causal links.

**Fig. 4.**
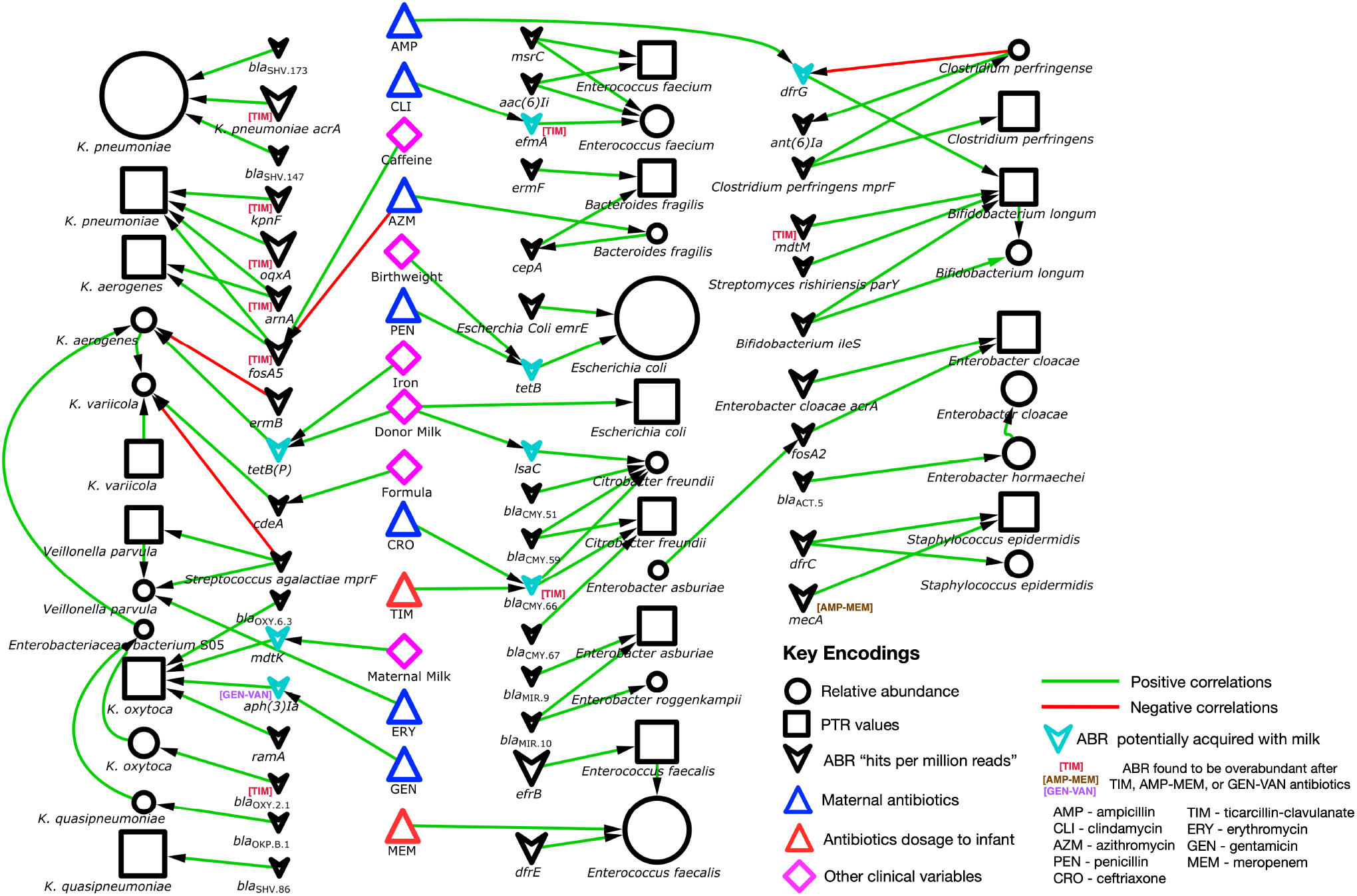
Causal network for infant gut microbiome. Node sizes are proportional to the average value.

#### Predicting provenance of ABR genes

An edge directed from ABR gene to taxon abundance was surmised to mean that the taxon was the predicted source for that ABR gene. We found 48 such edges (see Table 2), of which 25 were validated as described below. In general, because of horizontal gene transfers, it is often impossible to precisely infer the an organism carrying the ABR gene. The CARD database [39] provides prevalence values giving the percentage of whole-genome shotgun assemblies of the taxon from the NCBI repository that have at least one hit to the predicted ABR gene. The database also has annotations to original studies that indicate the species from which a given gene was originally isolated. Validated genes were present in at least 50% of the assemblies or were first isolated from the predicted species.

**Table 2.**
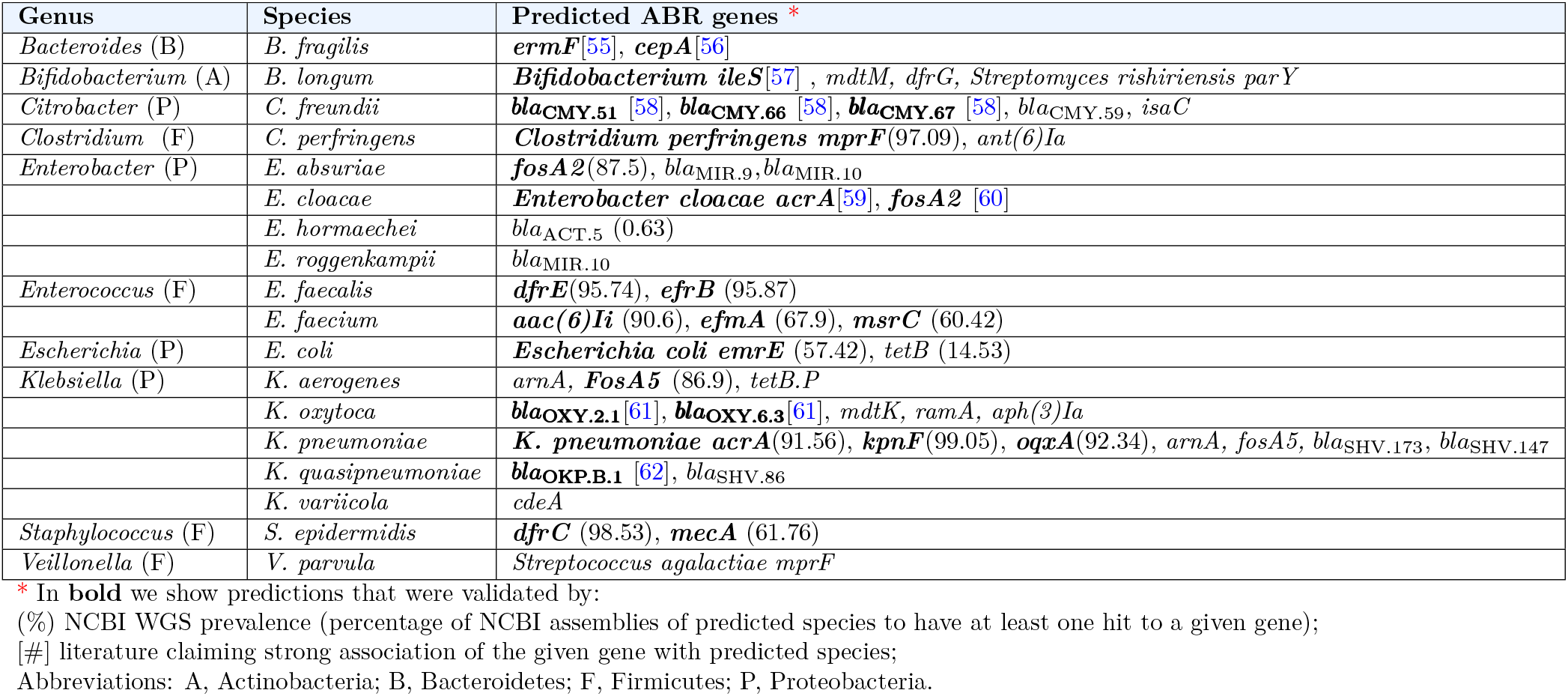
Predicted provenance of ABR genes from causal network

#### Human milk influence infant gut resistome

Human milk (either maternal or donor) seems to have an influence on ABR genes in the infant gut microbiota. Though, we lack the mothers microbiome data, it is not far-fetched to suggest the human milk may be the source of ABR genes. The presence of ABR genes in cow milk has been demonstrated [74]. Edges out of blue triangle nodes (corresponding to antibiotics administered to the mother) suggest a clear influence on taxa abundance and ABR genes in the infant gut.

#### Impact of antibiotics on bacterial taxa

Surprisingly, antibiotic dosage shows little effect on microbial taxa. Only a few edges are coming from variables associated with antibiotic dosage administered to infants (red triangles in Fig. 4). This fact can be the implication of the model construction assumptions and restriction rules. For example, we include only major bacterial taxa that are present in at least 5% of the samples with read coverage at least 5x. Therefore, included species are likely to be antibiotic-resistant, which explains the lack of direct impact of antibiotics. Besides, factors such as data sparsity, high dimensionality, errors in metagenomic sequencing and read mapping, and hidden confounders contribute to reduction in model sensitivity.

Nevertheless, the causal graph suggests that MEM causes an increase in abundance of *Enterococcus faecalis*. *Enterococcus faecalis* is a known early gut colonizer [75]. Enterocci are also a leading cause of hospital-acquired infections [76]. MEM is not effective against this pathogen [77]. However, antibiotic treatment can contribute to increase in the enteroccci abundance leading to reduction in the protective commensals and increase in *Clostridium* spp. [78].

A path from TIM → *bla*_CMY.66_ → *Citrobacter* (both PTR and abundance), suggests that *Citrobacter freundii* express *bla*_CMY.66_ encoding *β*-lactamases that inactivate TIM. The presence of this gene in *Citrobacter freundii* has been previously shownn [79]. This finidng argues that without knowing *apriori* the source of a gene, an educational guess could be made through our proposed novel approach. The presence of ABR strain of *Citrobacter freundii* is of major concern among pediatric population as it has been known to caused serious neonatal infections [80].

#### Impact of ABR genes on replication and abundance

The causal analysis sheds light on which ABR genes played the most critical role in the survival of some pathogens against antibiotic stress. Even though, according to the causal network, antibiotics dosage directly affects only a few ABR genes, we still can identify those that were most active against a particular drug. Using a Mann-Whitney U test, we identified genes that were significantly abundant after specific treatment. Such genes are indicated in colored font within square brackets in Fig. 4.

For example, *bla*_OXY.2.1_ is annotated with [TIM] meaning that number of hits to the sequence were significantly higher in infants with TIM administered within 48 hours prior to sampling. Edge leading from the gene to abundance of *K. oxytoca* suggest its role in allowing this pathogen to thrive.

Most importantly, our analysis suggests how *K. pneumonia* becomes dominant and actively replicating after the administration of TIM antibiotics. Fig. 4 indicates that five of ABR genes overabundant in TIM group have direct edges to PTR or abundance of this species. In particular, we hypothesize that *fosA5, oqxA, kpnF, arnA*, and *acrA* are possibly contributing to an increase in growth of *K. pneumonia*.

ABR genes are particularly influential when edges to both the abundance and PTR nodes of bacterial taxa. For example, difC helps *Staphylococcus epidermidis* to grow in presence of AMP and MEM.

A previous study [21] found that MFS genes are associated with increased replication for any bacterial taxa. In our study, we show that the influence of ABR genes on bacterial growth is not bound to specific gene families, but on the species that carries it and the type of antibiotics. Causal links between an ABR gene and a PTR node does not necessarily imply direct involvement in increasing regulation, which may be the result of nutrients availability or other indirect causes.

## 5. Conclusion

In this work, we propose a new computational pipeline called PeTRi, the first software package available for distributed computation of bacterial replication rates (PTR), which uses a reference-based approach with all available complete microbial genomes from the RefSeq database. Our approach shows how the replication rates computed by PeTRi can be used in the construction of a model for causal structural learning to infer potential cause–effect relationships between bacterial replication, bacterial abundance, and environmental factors.

When integrated into any whole metagenome sequence analysis, we obtain relative abundance values along with replication rates of the bacteria in the microbiome. Even though replication rate data is readily available, it is not usually computed. Here we apply this enhanced analysis pipeline on a publicly available infant gut microbiome dataset, and show that it can shed light on microbiome dynamics and provide new insights on antibiotic resistance.

Our results confirm that in general, higher replication does not imply growth in abundance. In fact, taxa with high abundance on average have low replication rates (i.e., in stationary phase).

Replication rates are found to be lower after the administration of *β*-lactams like ampicillin and meropenem, perhaps because they target dividing cells. However, some pathogens were actively replicating in the presence of *β*-lactam compound, TIM. In fact, pathogen *K. pneumoniae* was the most abundant and actively growing after the administration of this antibiotic. *K. pneumoniae* has been shown to be pathogenic and able to increase the mortality rate in preterm infants by 30 percent [81]. The causal network suggests that genes *fosA5, oqxA, kpnF, acrA,* and *arnA* had a causal effect on the abundance and/or replication rate. Therefore, *K. pneumoniae* has the necessary resistome to thrive after TIM treatment, and this drug should be avoided for premature infants.

Other interesting conclusions from the causal analysis include the influence of mother or donor milk on the resistome present in the infant’s gut. Thus, human milk should be tested for the presence of ABR genes [82].

Our methods show that even if an antibiotic resistance gene is novel, or in the absence of the CARD database, our method is still likely to identify its role in the microbiome.

The results of causal analysis need to be tested in the laboratory. Errors in inference may be caused by small sample size, errors in sequencing, errors in mapping reads to the correct genomes, errors in abundance matrix, presence of undocumented or new bacterial taxa, inability to measure known or hidden confounders related to the diet and environment of the mother and infant, and much more. Nevertheless, we set the stage for a novel approach to study microbiome dynamics from studies with limited time points.

## Acknowledgements

The authors thank the members of the Bioinformatics Research Group (BioRG) at FIU for valuable feedback and comments.

